# Paranoid schizophrenia and methamphetamine-induced paranoia are both characterized by a similar LINE-1 partial methylation profile, which is more pronounced in paranoid schizophrenia

**DOI:** 10.1101/403535

**Authors:** Rasmon Kalayasiri, Korakot Kraijak, Apiwat Mutirangura, Michael Maes

## Abstract

**Background:** There is evidence that schizophrenia is a neuro-immune disorder. Genes linked to intragenic LINE-1 methylation show a strong association with immune-associated disorders including psychosis. The aim of this study was to examine LINE-1 methylation patterns in paranoid schizophrenia and methamphetamine-induced paranoia, a model for schizophrenia.

**Methods:** This study recruited 31 patients with paranoid schizophrenia, 94 with methamphetamine-induced paranoia (MIP) and 163 normal controls. LINE-1 methylation patterns were assayed in peripheral blood mononuclear cells and a combined bisulphite restriction analysis and COBRA were used to estimate global methylation (mC) and LINE-1 CpG dinucleotide methylation patterns, namely 2 methylated (mCmC) and 2 unmethylated (uCuC) CpGs and the partially methylated loci mCuC (5’m with 3’u) and uCmC (5’u with 3’m).

**Results:** Patients with paranoid schizophrenia show highly significant changes in LINE-1 partial methylation patterns, namely a higher % mCuC and lower % uCmC as compared with controls and MIP patients, while the latter show higher % mCuC but lower % uCmC as compared with controls. Higher % mCuC significantly predicts paranoid schizophrenia with a sensitivity of 51.6%, specificity of 97.5% and an area under the ROC curve of 0.895.

**Conclusions:** The results indicate that a common dysfunction in LINE-1 partial methylation may underpin both paranoid schizophrenia and MIP and that this methylation pattern is significantly more expressed in paranoid schizophrenia than MIP. Reciprocal links between impairments in LINE-1 methylation and neuro-immune and neuro-oxidative pathways may underpin the pathophysiology of both MIP and paranoid schizophrenia.

## Introduction

There is now evidence that schizophrenia is characterized by immune-inflammatory alterations, including activated immune-inflammatory responses, a subchronic mild inflammatory response and activated nitro-oxidative processes (Anderson et al., 2013; Davis et al., 2016; Smith & Maes, 1995). Major findings indicate signs of an acute phase response (e.g. increased haptoglobin and complement factors), M1 macrophagic activation (e.g. increased production of interleukin (IL)-1β, IL-6, and tumor necrosis factor-α), T helper (Th)-1, Th-2 and Th-17 responses (e.g. increased levels of IL-2, IL-17, IL-4, IL-5, and interferon-γ), a T regulatory (Treg) response (e.g. increased IL-10 levels) and indicants of oxidative stress including lowered lipid-associated antioxidant defenses and increased lipid peroxidation (Brinholi et al., 2015; Maes, Meltzer, & Bosmans, 1994; Noto, Maes, et al., 2015; Noto, Ota, et al., 2015; Noto et al., 2014; Noto et al., 2016; Zeni-Graiff et al., 2016). Many M1 macrophagic and Th-1 products have detrimental effects on the brain (Leonard & Maes, 2012) collectively named neuroprogression, which includes aberrations in neuronal plasticity, dendrite growth neurogenesis, apoptosis, receptor expression and functioning, including the NMDA receptors, and thus changes in brain neuro-circuitry (Anderson & Maes, 2013; Smith & Maes, 1995). Schizophrenia is also accompanied by changes in leukocyte telomere length with for example a lowered telomere length in individuals at high risk for psychosis (Maurya et al., 2018; Maurya et al., 2017). There are also data that schizophrenia is accompanied by changes in DNA methylation patterns although a recent systematic review showed inconsistent findings, including hypermethylation, hypomethylation or no changes in global methylation patterns (Teroganova, Girshkin, Suter, & Green, 2016).

Most of the DNA methylation occurs in regions with a high percentage of CpG sites that are concentrated in promoter regions, namely the 5’-CpG-3’ islands and this methylation process may be accompanied by gene silencing (Bird, 2002; Kitkumthorn & Mutirangura, 2011). Many studies measured cytosine methylation of LINE-1 (long interspersed element-1s), which are interspersed repetitive sequences and retrotransposons (Kitkumthorn & Mutirangura, 2011). LINE-1 methylation patterns play a key role in maintaining genomic integrity whilst intragenic LINE-1 mediates gene expression in cis (Kemp & Longworth, 2015; Kitkumthorn & Mutirangura, 2011; Wangsri, Subbalekha, Kitkumthorn, & Mutirangura, 2012). LINE-1 hypomethylation may cause repression of gene expression, genomic instability, and aberrations in DNA repair genes and DNA double-strand break repair (Aporntewan et al., 2011; Pornthanakasem et al., 2008; Wangsri et al., 2012). As such, hypomethylation of promotor LINE-1 retrotransposable elements is frequently associated with human disorders including cancer (Chalitchagorn et al., 2004; Dammann et al., 2005). Most importantly, genes associated with the cis-regulatory activities of intragenic LINE-1 show an association with immune and oxidative pathways, apoptosis and cell differentiation and many (auto)immune and neuroprogressive diseases, possibly including schizophrenia (Wanichnopparat, Suwanwongse, Pin-On, Aporntewan, & Mutirangura, 2013).

Recently, we have shown that the assay of LINE-1 methylation patterns with the combined bisulphite restriction analysis of LINE-1s (COBRALINE-1) methylation is superior as compared with measurements of global methylation (Kitkumthorn, Keelawat, Rattanatanyong, & Mutirangura, 2012; Kitkumthorn, Tuangsintanakul, Rattanatanyong, Tiwawech, & Mutirangura, 2012). This method allows to measure LINE-1 overall methylation levels (indicating global DNA methylation) as well as to classify LINE-1 alleles into 4 patterns based on the methylation status of two CpG dinucleotides on each strand from 5’ to 3’, namely 2 methylated (mCmC) and 2 unmethylated (uCuC) CpGs and two types of partially methylated loci (mCuC that is 5’m with 3’u and uCmC that is 5’u with 3’m CpGs) (Wangsri et al., 2012). For example, in peripheral blood mononuclear cells (PBMCs), the uCuC LINE-1 hypomethylated loci show a significantly better diagnostic performance for oral cancer than overall methylation (Kitkumthorn, Tuangsintanakul, et al., 2012). Nevertheless, there are no data on COBRALINE-1 assays in schizophrenia.

Psychosis including paranoia and schizophrenia-like cognitive deficits are frequently induced by use of methamphetamine (streetnames: meth, ice, crystal, yaba), a psychostimulant substance (Hsieh, Stein, & Howells, 2014; Kalayasiri, Mutirangura, Verachai, Gelernter, & Malison, 2009; Kalayasiri, Verachai, Gelernter, Mutirangura, & Malison, 2014). Such effects are ascribed to enhanced release and turnover of dopamine and glutamate in the brain with ensuing damage to cortical GABA-ergic neurons (Hsieh et al., 2014; Vasan & Olango, 2018). Nevertheless, methamphetamine may activate immune-inflammatory and oxidative pathways, for example by effects on DNA oxidation, the anti-cholinergic anti-inflammatory pathway and gut microbiota causing increased gut permeability (Prakash et al., 2017; Ramkissoon & Wells, 2015). Such effects may at least in part determine MA-induced neuroprogressive effects including glutamate neurotoxicity, damage to neuronal dendrites, neuronal death in frontal, prefrontal and temporal lobes, white matter gliosis and hypertrophy (Prakash et al., 2017; Thompson et al., 2004), which in turn, may cause methamphetamine-induced paranoia (MIP) and deficits in information processing, episodic memory and executive functions (Hsieh et al., 2014; Prakash et al., 2017; Thompson et al., 2004). Interestingly, MIP is considered to be a model for schizophrenia (Hsieh et al., 2014; Shelly et al., 2016). Nevertheless, no studies have directly compared LINE-1 methylation profiles between paranoid schizophrenia and MIP.

Hence, the aim of the present study is to examine COBRALINE-1-derived methylation profiles between patients with paranoid schizophrenia and MIP and normal controls. Similar changes in partial LINE-1 methylation patterns in both MIP and paranoid schizophrenia may point toward a common pathophysiology related to aberrations in LINE-1 methylation.

## Subjects and methods

### Participants

We recruited subject with MIP who were admitted to the Princess Mother National Institute on Drug Abuse Treatment (PMNIDAT) and patients with paranoid schizophrenia who were treated at the Department of Psychiatry, King Chulalongkorn Memorial Hospital, Bangkok, Thailand. Normal controls were recruited by word of mouth at the blood donation center, Thai Red Cross Society, Bangkok, Thailand. All participants were Thai nationals of both genders and 18-65 years old. The diagnostic assessments of substance use were made employing the Thai version of the Semi-Structured Assessment for Drug Dependence and Alcoholism (SSADDA), while the diagnosis of schizophrenia, paranoid subtype, and MIP were made using the Diagnostic and Statistical Manual of Mental Disorders (DSM-IV) (American Psychiatric Association, 2000) and Methamphetamine Experience Questionnaire (MEQ) – Thai version (Kalayasiri et al., 2014) by exploring paranoid experiences during MA use. We have excluded paranoid schizophrenia patients who were diagnosed with substance use disorder, while we excluded MIP subjects who were diagnosed with primary psychotic disorders or schizophrenia. Moreover, we excluded patients and controls with a lifetime history of bipolar disorder, major depressive disorder, psycho-organic disorders; neurodegenerative / neuroinflammatory disorders (e.g. Parkinson’s disease, multiple sclerosis and stroke), neurologic disorders (e.g. epilepsy and brain trauma) and (auto)immune disorders (e.g. systemic lupus erythematosus). The study was approved by the Human Ethics Committee of the Faculty of Medicine, Chulalongkorn University (Med Chula IRB #417/57).

### Measurements

Socio-demographic data and substance use variables and related diagnoses were obtained by trained clinical psychologists certified for SSADDA interview employing the SSADDA. The principal investigator (R.K.) made the diagnoses of MIP and paranoid schizophrenia using the SSADDA, MEQ and DSM-IV-TR criteria for paranoid schizophrenia. The MEQ shows a good inter-instrument reliability (K = 0.87) (Kalayasiri et al., 2014). The body mass index (BMI) was computed as body weight (kg) / length (meter)^2^. Current smoking (last 6 months) and use of psychotropic medications was registered as dummy variables (yes/no).

### DNA extraction, bisulfite modification, and COBRALINE-1s

DNA extraction and COBRALINE-1 assays were carried out at the Center for Excellence in Molecular Genetics of Cancer and Human Diseases, Faculty of Medicine, Chulalongkorn University, Bangkok, Thailand. Blood samples were centrifuged at 1,000 g for 10 minutes to collect PBMCs and stored at −80 °C until assayed for COBRALINE-1 assays. Whole blood was extracted by 1) adding lysis buffer with 10% sodium dodecyl sulfate and proteinase K and then incubated overnight at 50 ˚C, 2) purifying using phenol-chloroform and centrifuging at 4 ˚C with 14,000 g for 15 minutes, 3) precipitating the DNA pellet using 10 Molar ammonium acetate and absolute ethanol, 4) washing DNA pellet by 70% ethanol, and 5) drying DNA pellet and dissolving it by Tris-EDTA.

The combined bisulphite restriction analysis of LINE-1s (COBRALINE-1) was used for determining pattern of LINE-1s methylation. A total of 1 microgram (μg) of DNA was used in the bisulfite treatments that converted unmethylated cytosine to uracil, while methylated cytosine was not changed. The bisulphite DNA modification was performed using the EZ-DNA methylation kit and specific primers is LINE-1s-F (5’GTTAAAGAAAGGGGTGA YGGT-3’) and LINE-1s-R (5’AATACRCCRTTTCTTAAACC RATCTA-3’) at 95 ˚C denature for 15 minutes, 50 ˚C annealing for 35 cycles and 72 ˚C final extension. LINE-1s were digested with *Taq*I and *Tas*I at 65 ˚C overnight. The digested products of bisulfite-treated LINE-1s were separated to strands with different length including 92 (^m^C^u^C), 60 (^u^C^u^C), 50 (^m^C^m^C), 42 (^m^C^m^C and ^u^C^m^C), and 32 (^u^C^u^C and ^u^C^m^C) base pairs (bp) that were measured by using polyacrylamide gel electrophoresis and stained with SYBR. We used deionized water as a negative control and HeLa, Daudu, and Jurkat as positive control.

The intensity of each band was assigned into A, B, C, D, E (e.g., A = %92/92, B = %60/56, C = %50/48, D = %42/40, E = %32/28). The intensity of 18 bp was calculated and assigned to F = ((D + E) – (B + C))/2). Percentage of each patterns of DNA methylation were calculated by using the following formula:

% overall methylation = ((A + 2C + F)*100) / (2A + 2B + 2C + 2F)

% (^m^C^m^C) hypermethylation = ((C/2)*100) / ((C/2) + A + B + F)

% (^u^C^m^C) partial methylation = (F*100) / ((C/2) + A + B + F)

% (^m^C^u^C) partial methylation = (A*100) / ((C/2) + A + B + F)

% (^u^C^u^C) hypomethylation = (B*100) / ((C/2) + A + B + F)

### Statistical Analyses

We used analysis of contingency tables (X^2^-tests) to check associations between categorical variables and used analysis of variance (ANOVA) to check differences in scale variables between diagnostic groups. We used multivariate general linear model (GLM) analysis to examine the associations between diagnosis (three groups, namely paranoid schizophrenia, MIP and controls) and LINE-1 methylation patterns while adjusting for sex, age and BMI. Consequently, we used tests for between-subject effects to assess the associations between diagnosis and the separate LINE-1 methylation data. Model-generated estimated marginal mean values were computed and protected post-hoc analyses were used to assess pairwise differences between categories. We used binary logistic regression analysis to define the most important predictors of paranoid schizophrenia versus controls, paranoid schizophrenia versus MIP and paranoid schizophrenia versus MIP+controls. Odds ratios and 95% confidence intervals were computed, while Nagelkerke values were used to estimate effect size. We used the area under the Receiver Operating Curve (ROC) coupled with sensitivity and specificity to estimate the overall diagnostic performance. In addition, we computed 2 different z weighted composite scores based on the partial LINE-1 methylation data, namely a) sum of the two partial methylation data as z transformation of % mCuC (z mCuC) + z uCmC (reflecting overall partial LINE-1 methylation), and b) z mCuC – z uCmC (reflecting the shift among these partial methylation profiles). All statistical analyses were performed using IBM SPSS windows version 24. Tests were 2-tailed and an alpha level of 0.05 indicated statistically significant results.

## Results

**Table 1** shows the socio-demographic and biomarker data in paranoid schizophrenia, MIP and controls. There were no significant differences in age and BMI between the study groups, while there were more males in the MIP group as compared with controls and paranoid schizophrenia. This table also shows the estimated-marginal mean values of the LINE-1 methylation data in the three diagnostic groups after adjusting for age, sex and BMI. **Figure 1** shows the LINE-1 methylation profile displayed as z values (mean=0 and standard deviation=1) in the three diagnostic groups. **Table 2** shows the results of multivariate GLM analysis with 7 DNA methylation data as dependent variables and diagnosis (three groups) as primary explanatory variable while adjusting for age, sex and BMI. There was a highly significant effect of diagnosis with a partial eta squared = 0.151, after considering the effects of age, sex and BMI, which were not significant. Table 2 shows the results of the tests for between-subject effects and protected post-hoc analysis. There was a strong association between diagnosis and % mCuC (partial eta squared = 0.217). % mCuC was significantly different between the three groups and increased from controls → MIP → paranoid schizophrenia. There were also significant differences in % uCmC between the groups, although the effect size was smaller (partial eta squared = 0.056). % uCmC decreased from controls → MIP → paranoid schizophrenia. There were also significant differences in the z composite scores z mCuC + z uCmC (effects size = 0.051) and z mCuC – z uCmC (effect size = 0.165). The z mCuC + z uCmC score was significantly higher in both patient groups as compared with controls, while the z mCuC – z uCmC score was significantly different between the three groups and increased from controls → MIP → paranoid schizophrenia.

**Table 1.** Demographic data and model-generated estimated marginal mean values of LINE-1 methylation data in healthy controls (HC), Patients with methamphetamine-induced paranoia (MIP) and patients with schizophrenia, paranoid subtype (SCZ).

| Variables | HC <sup>A</sup> | MIP <sup>B</sup> | SCZ <sup>C</sup> | F / $\chi^2$ | df | p |
| --- | --- | --- | --- | --- | --- | --- |
| Age (years) | 33.8 (11.6) | 33.0 (6.2) | 37.2 (9.6) | 2.11 | 2/285 | 0.122 |
| Sex (Female/male) | 82/81 <sup>B</sup> | 31/63 <sup>A,C</sup> | 18/13 <sup>B</sup> | 9.44 | 2 | 0.009 |
| BMI (kg/m<sup>2</sup>) | 23.9 (4.3) | 25.0 (2.2) | 23.8 (4.3) | 2.88 | 2/285 | 0.058 |
| % overall mC | 67.1 (0.2) | 66.6 (0.3) | 66.5 (0.5) | 1.25 | 1/282 | 0.288 |
| % mCmC | 33.9 (0.3) | 33.3 (0.4) | 33.3 (0.6) | 1.06 | 1/282 | 0.349 |
| % uCmC | 32.3 (0.1) <sup>B,C</sup> | 31.8 (0.2) <sup>A,C</sup> | 31.0 (0.3) <sup>A,B</sup> | 8.30 | 1/282 | <0.001 |
| % mCuC | 18.6 (0.2) <sup>B,C</sup> | 20.3 (0.2) <sup>A,C</sup> | 21.9 (0.4) <sup>A,B</sup> | 39.01 | 1/282 | <0.001 |
| % uCuC | 21.9 (0.2) | 22.0 (0.3) | 22.4 (0.2) | 0.63 | 1/282 | 0.631 |
| z mCuC + z uCmC | -0.193 (0.077) <sup>B,C</sup> | 0.186 (0.104) <sup>A</sup> | 0.419 (0.177) <sup>A</sup> | 7.65 | 1/282 | 0.001 |
| z mCuC – z uCmC | -0.322 (0.072) <sup>B,C</sup> | 0.244 (0.098) <sup>A,C</sup> | 0.895 (0.166) <sup>A,B</sup> | 27.96 | 1/282 | <0.001 |

Results are shown as mean (SD)
<sup>A,B,C</sup>: significant differences between group means
BMI: body mass index

**Table 2.** Associations between diagnostic groups, healthy controls (HC), paranoid schizophrenia (SCZ) and methamphetamine-induced paranoia (MIP) and LINE-1s methylation, while adjusting for background variables

| Multivariate Tests | Dependent variables | Explanatory variables | F | df | p |
| --- | --- | --- | --- | --- | --- |
| #1 | All 7 methylation data | HC / MIP / SCZ | 10.12 | 10 / 558 | <0.001 |
|  |  | Age | 0.26 | 5 / 278 | 0.936 |
|  |  | Female / Male | 1.33 | 5 / 278 | 0.253 |
|  |  | BMI | 1.43 | 5 / 278 | 0.213 |
| #2 | All 7 methylation data | Smoking | 1.68 | 5 / 215 | 0.141 |
| #3 | All 7 methylation data | Neuroleptics | 0.07 | 5 / 110 | 0.997 |
| #4 | All 7 methylation data | Atypical antipsychotics | 0.31 | 5 / 110 | 0.905 |
| #5 | All 7 methylation data | Antidepressants | 0.51 | 5 / 110 | 0.765 |
| #6 | All 7 methylation data | Benzodiazepines | 0.47 | 5 / 110 | 0.799 |

HC / MIP / SCZ: healthy controls, methamphetamine-induced paranoia and schizophrenia, paranoia subtype
BMI: body mass index
7 methylation data: entered are % overall mC, % mCmC, % uCmC, % mCuC, % uCuC, z uCmC + z mCuC, and z mCuC - z uCmC

**Figure 1.**
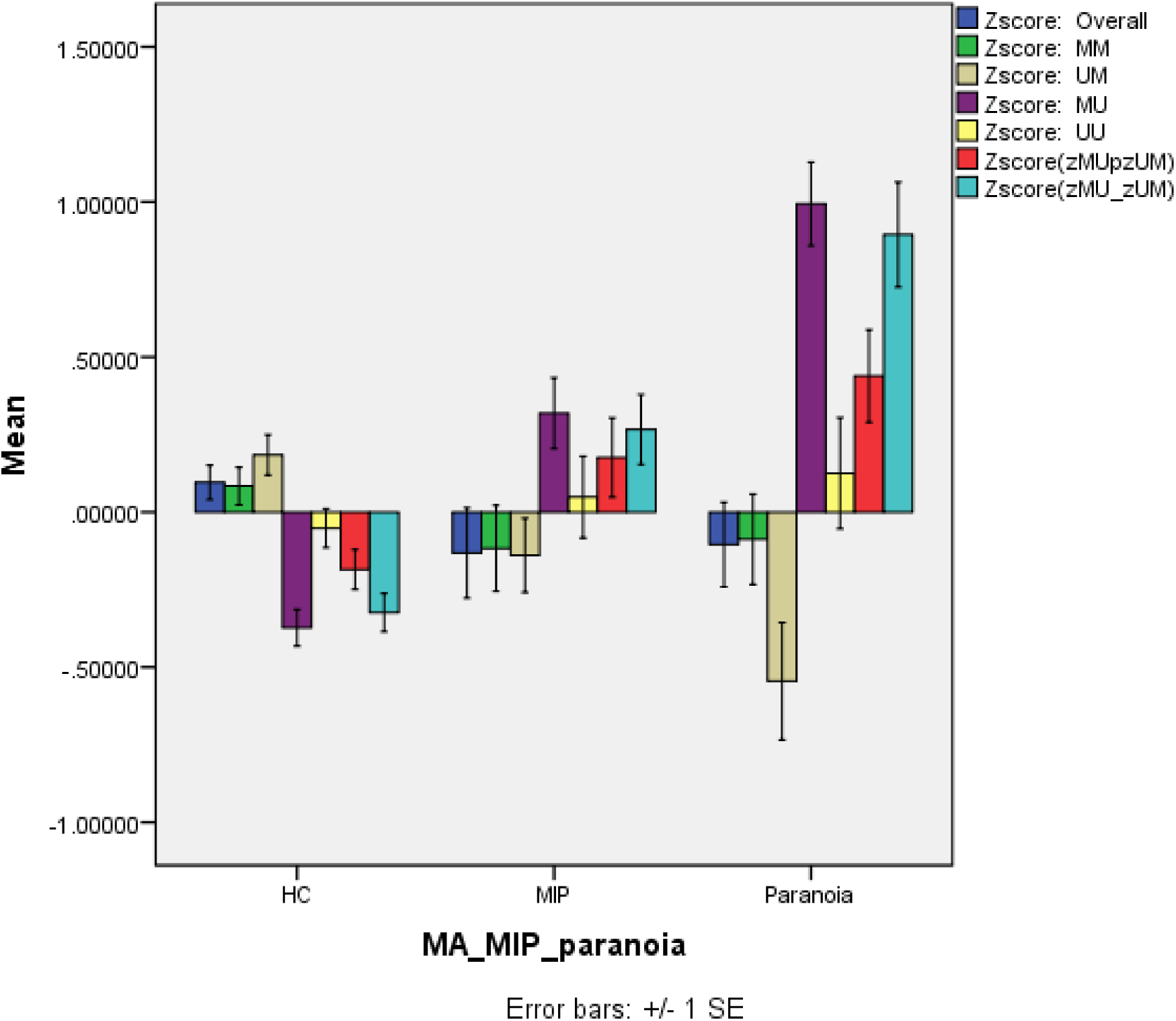
LINE-1 methylation profile displayed as z values (mean=0 and standard deviation=1) in healthy controls (HC), methamphetamine-induced paranoia (MIP) and paranoid schizophrenia (paranoia). MM/UU: 2 methylated (mCmC; MM) and 2 unmethylated (uCuC; UU) CpGs. UM/MU: partially methylated loci mCuC (5’m with 3’u; MU) and uCmC (5’u with 3’m; UM). zMUpzUM: computed as z transformations of % mCuC (z mCuC) + z uCmC (reflecting overall partial LINE-1 methylation). zMU_zUM: computed as z mCuC – z uCmC (reflecting the shift among these partial methylation profiles).

We have also examined whether there were any effects of smoking and drug use on the LINE-1 methylation data by entering these data in GLM analysis. Table 2 shows the results of these GLM analyses. There were no significant effects of smoking (versus not smoking), and use of neuroleptics, atypical antipsychotics, antidepressants and benzodiazepines on the results.

**Table 3** shows the results of binary logistic regression analyses with paranoid schizophrenia or paranoid schizophrenia + MIP (any psychosis) as dependent variables and the LINE-1 methylation data (alone or combined with sex, age and BMI) as independent variables. We found (regression #1) that increased % mCuC significantly predicted paranoid schizophrenia versus controls with a Nagelkerke value of 0.513. The area under the ROC was 0.895 and 90.2% of all cases were correctly classified with a sensitivity = 51.6% and specificity = 97.5%. Regression #2 shows that any psychosis was best predicted by % mCuC with an effect size = 0.246. The area under the ROC was 0.740 and 70.5% of all cases were correctly classified with a sensitivity = 59.3% and specificity = 79.1%. % mCuC was also significant in discriminating paranoid schizophrenia from MIP and from MIP + controls.

**Table 3.** Results of binary logistic regression analyses with healthy controls (HC), paranoid schizophrenia (SCZ), and methamphetamine-induced paranoia (MIP) as dependent variables and LINE-1 methylation patterns, demographic and clinical data as explanatory variables.

| Dichotomy | Explanatory variables | Wald | df | p | OR | 95% CI | Nagelkerke ROC curve (SE) |
| --- | --- | --- | --- | --- | --- | --- | --- |
| #1. SCZ versus HC | % mCuC | 33.53 | 1 | <0.001 | 12.28 | 5.26 – 28.71 | Nagelkerke: 0.513<br>ROC: 0.895 (0.030) |
| #2. SCZ + MIP versus controls | % mCuC | 43.65 | 1 | <0.001 | 2.93 | 2.13 – 4.04 | Nagelkerke: 0.246<br>ROC: 0.740 (0.030) |
| #3. SCZ versus MIP | % mCuC | 8.67 | 1 | 0.003 | 2.03 | 1.26 – 3.26 | Nagelkerke: 0.314<br>ROC: 0.720 (0.049) |
|  | Sex | 10.03 | 1 | 0.002 | 0.19 | 0.07 – 0.53 |  |
|  | Age | 4.96 | 1 | 0.026 | 1.07 | 1.01 – 1.14 |  |
|  | BMI | 6.72 | 1 | 0.010 | 0.81 | 0.70 – 0.95 |  |
| #4. SCZ versus HC + MIP | % mCuC | 27.03 | 1 | <0.001 | 3.39 | 2.14 – 5.37 | Nagelkerke: 0.274<br>ROC: 0.831 (0.034) |
|  | Sex | 5.41 | 1 | 0.020 | 0.36 | 0.15 – 0.85 |  |
|  | Age | 4.41 | 1 | 0.036 | 1.05 | 1.03 – 1.09 |  |

OR: Odd's ratio
95% CI: 95% confidence interval, lower and upper limit

## Discussion

The first major finding of this study is that PBMCs of paranoid schizophrenia patients show a higher % mCuC and lowered % uCmC DNA partial methylation as compared with controls and that the increase in % mCuC partial methylation was predictive for paranoid schizophrenia with an area under the ROC of 0.895. These findings show that paranoid schizophrenia is accompanied by aberrations in LINE-1 partial methylation, while there are no significant changes in global methylation and no signs of hypermethylation or hypomethylation. These data show that classifying LINE-1 alleles into four classes based on the methylation status of 2 CpG dinucleotides, including partial methylation profiles, provides more specific information than the assessment of global LINE-1 methylation alone. Previously, it was reported that the COBROLINE-1 patterns (namely uCuC hypomethylated LINE-1 loci) assayed in PBMCs better discriminate patients with oral cancer from normal controls as compared with global methylation data (Kitkumthorn, Keelawat, et al., 2012). Phrased differently, if we had assayed only global LINE-1 methylation we would have obtained negative results, while our COBRALINE-1 analyses show a highly significant association with partial methylation patterns. Our results are difficult to compare with previous DNA and LINE-1 methylation data in schizophrenia because we assayed a more complete LINE-1 methylation pattern, while previous studies in schizophrenia measured global methylation. Likewise, a systematic review on DNA methylation in schizophrenia (Teroganova et al., 2016) reported very inconsistent results pointing towards hypo- or hypermethylation or no changes at all.

The second major finding of this study is that both paranoid schizophrenia and MIP show a similar LINE-1 methylation profile, namely increased % mCuC and lowered % uCmC partial LINE-1 methylation compared with normal controls, while these changes are significantly more pronounced in paranoid schizophrenia as compared with MIP. This is important as MIP is considered to be a model of schizophrenia (Hsieh et al., 2014). Indeed, methamphetamine use is frequently accompanied by paranoid symptoms and cognitive impairments, including episodic memory and executive functions, which are quite similar with those observed in paranoid schizophrenia (Glasner-Edwards & Mooney, 2014; Hsieh et al., 2014; Prakash et al., 2017; Thompson et al., 2004). The latter authors ascertain that the differential diagnosis of primary schizophrenia versus MIP may sometimes be challenging although MIP is frequently a transient condition and defined by a temporal relationship between methamphetamine use and the consequent psychosis, while s“schizophrenia is not related to drug intoxication or withdrawal” (Glasner-Edwards & Mooney, 2014; World Health Organization, 1992). In another study, we established that use of methamphetamine increases % mCuC LINE-1 methylation but does not change % uCmC LINE-1 methylation (Kalayasiri et al., submitted), while here we showed that MIP is accompanied by significantly lowered % uCmC. Phrased differently, aberrations in both partial methylation patterns and a shift from lowered % uCmC towards increased % mCuC is a characteristic of paranoia, either drug-induced or primary paranoia. If aberrations in LINE-1 methylation patterns are causally linked to psychosis, we may hypothsize that a) methamphetamine may increase the vulnerability to paranoia by increasing % mCuC partial methylation; and b) the greater dysfunctions in LINE-1 methylation in paranoid schizophrenia versus MIP could reflect differences in severity of psychosis with a worse phenomenology in paranoid schizophrenia than MIP.

Previously, it was proposed that an increased release of central dopamine and glutamate and damage to GABA-ergic neurons in the cortex could underpin both MIP and paranoid schizophrenia (Hsieh et al., 2014; Vasan & Olango, 2018). Nevertheless, here we show that an epigenetic process may constitute a common pathophysiology underpinning both types of paranoid psychoses. There is now some evidence that reactive oxygen species may induce changes in LINE-1 methylation. For example, in gold miners, oxidative stress associated with increased mercury levels, is associated with increased LINE-1 methylation (Narvaez et al., 2017). LINE-1 methylation is reduced in cells treated with peroxides (Kloypan, Srisa-art, Mutirangura, & Boonla, 2015), while antioxidants, including alpha-tocopherol, N-acetyl cysteine and NAC, S-adenosylmethionine and folic acid, may restore LINE-1 hypomethylation. In UM-UC-3 bladder cell (carcinoma) lines, peroxides induced LINE-1 hypomethylation which could be restored by alpha-tocopherol, while treatment with peroxides increased methylation of the runt-related transcription factor 3 (RUNX3) promotor, a tumor suppressor gene, which was attenuated by alpha-tocopherol (Wongpaiboonwattana, Tosukhowong, Dissayabutra, Mutirangura, & Boonla, 2013). Such results indicate that oxidative stress may promote carcinogenesis through modulation of LINE-1 methylation. Interestingly, in biliary atresia patients, LINE-1 hypomethylation is associated with signs of oxidative stress, while LINE-1 methylation is positively associated with relative telomere length (Udomsinprasert et al., 2016). In this respect, it is known that DNA methylation processes in subtelomeric DNA repeats are an important factor in the regulation of telomere length (Yehezkel, Segev, Viegas-Pequignot, Skorecki, & Selig, 2008). Moreover, inflammatory biomarkers (including ICAM-1) are negatively associated with leukocyte telomere length and global DNA methylation, which may increase genome instability (Dong et al., 2017). By inference, the neuro-immune and neuro-oxidative pathways that characterize schizophrenia and are inducible by methamphetamine could have affected LINE-1 methylation and consequently telomere length (see Introduction). There are, however, no data whether inflammatory and redox-related pathways may affect LINE-1 partial methylation processes. Importantly, genes linked with intragenic LINE-1 show a strong association with oxidative, inflammatory, immune and apoptotic processes as well as cell differentiation and with many autoimmune, immune, and neurodegenerative disorders, possibly also with schizophrenia (Wanichnopparat et al., 2013). These new epigenomic findings do not rule out that activated glutamate release and damage to GABA-ergic neurons in the brain may underpin MIP and paranoid schizophrenia (Hsieh et al., 2014). It was explained previously that activation of neuro-immune and neuro-oxidative pathways may cause increased glutamate toxicity leading to a number of neuronal dysfunctions including in GABA-ergic neurons (Morris et al., 2018; Morris, Carvalho, Anderson, Galecki, & Maes, 2016).

In conclusion, the results indicate that a shift in LINE-1 partial methylation patterns from % uCmC to increased % mCuC may underpin paranoid schizophrenia and MIP and that this pattern is significantly more pronounced in paranoid schizophrenia than MIP. Schizophrenia is now conceptualized as a neuro-immune and neuro-oxidative disorder, while methamphetamine may also induce these pathways. Induction of these paths may lead to alterations in LINE-1 methylation patterns, while genes linked with intragenic LINE-1 show an association with many neuro-immune disorders, including schizophrenia. Future research should examine the reciprocal links between neuro-immune and neuro-oxidative pathways and impairments in LINE-1 partial methylation in relation to the pathophysiology of both MIP and paranoid schizophrenia.

## Acknowledgement

We thank Mr. Prakasit Rattanatanyong and Dr Nakarin Kitkumthorn for excellent technical support in laboratory. We appreciate Ms Maturada Phetsung and Ms Sirapat Settayanon for the helps on laboratory work. We thank Drs Dolnapha Rattanakorn and Thitiporn Supasitthumrong for DNA samplings, Mr Wuthichai Hasook for healthy controls recruitment, and staffs at the Princess Mother National Institute on Drug Abuse Treatment for facilitating data collection.

## Contributors

All authors contributed to interpretation of the data and writing of the manuscript.

